# Single-cell transcriptional landscape of temporal neutrophil response to burn wound in larval zebrafish

**DOI:** 10.1101/2024.04.01.587641

**Authors:** Yiran Hou, Parth Khatri, Julie Rindy, Zachery Schultz, Anqi Gao, Zhili Chen, Angela LF Gibson, Anna Huttenlocher, Huy Q. Dinh

**Author notes:** Equally contributed (Yiran Hou, Parth Khatri). co-corresponding authors **Correspondence:** Anna Huttenlocher, Huy Q. Dinh.

## Abstract

Neutrophils accumulate early in tissue injury. However, the cellular and functional heterogeneity of neutrophils during homeostasis and in response to tissue damage remains unclear. Here, we use larval zebrafish to understand neutrophil responses to thermal injury. Single-cell transcriptional mapping of myeloid cells during a 3-day time course in burn and control larvae revealed distinct neutrophil subsets and their cell-cell interactions with macrophages across time and conditions. The trajectory formed by three zebrafish neutrophil subsets resembles human neutrophil maturation, with varying transition patterns between conditions. Through ligand-receptor cell-cell interaction analysis, we found neutrophils communicate more in burns in a pathway and temporal manner. Finally, we identified the correlation between zebrafish myeloid signatures and human burn severity, establishing GPR84+ neutrophils as a potential marker of early innate immune response in burns. This work builds the molecular foundation and a comparative single-cell genomic framework to identify neutrophil markers of tissue damage using model organisms.

## Introduction

Neutrophils are the first responders of the innate immune response to tissue damage and infection. As the most abundant circulating leukocyte, neutrophils are important drivers of these early inflammatory responses and can also play a role in chronic inflammation and tissue damage. Complex injuries, like burns, lead to robust innate immune inflammation and damaging tissue responses. Previous work has identified neutrophil activation in burn patients; however, the role of neutrophils is complicated and likely context-dependent (1, 2). In recent years, single-cell profiling of neutrophils in normal development and disease models has identified diverse neutrophil populations at the molecular level (3–6). Most studies have classified neutrophils based on their maturation status (7). Some studies further identified specific subsets of neutrophils that are positively or negatively associated with disease progression. One recent study focused on neutrophils in the early stage of severe burns and found human neutrophils matching the mouse G3/4/5a-c subsets in both burned and healthy donors (5, 8). These studies showcase the utility of single-cell transcriptomics to facilitate our understanding of neutrophil involvement in tissue damage contexts.

Although direct profiling of neutrophils from human tissues is feasible, there are limitations to what can be learned from human burns. Neutrophils are often isolated either from local burn wound tissue (eschar) or peripheral blood late after injury, which does not reflect the early state of neutrophils in tissues or provide information about neutrophil progenitors in the bone marrow (9). Zebrafish enable whole organism profiling of neutrophils after burns with unbiased sampling of neutrophils at all tissue locations including the hematopoietic tissue. Previous studies have demonstrated that zebrafish provide a robust model system to study thermal injury that enables the imaging of the temporal and spatial dynamics of the innate immune response to injury (10, 11). Recent scRNA-Seq also confirmed conserved maturation stages and markers of neutrophil subsets in zebrafish and humans (12), justifying the use of zebrafish as a model in studying human neutrophil responses to thermal injury. Using this model, we previously found that both human burn and zebrafish burn tissue showed an increase in IL6 production (11). In zebrafish, we found that the IL6 receptor was important for neutrophil but not macrophage recruitment to burn wounds, however the mechanism for these effects remains unclear.

Here, we used transcriptional profiling at single cell resolution of neutrophils and macrophages in larval zebrafish to compare gene expression during homeostasis and following a burn wound. We describe the molecular diversity within neutrophils and macrophages during the early innate immune response to burn. By comparing between burned and unwounded conditions, we show subset-level functional changes and subset-to-subset transitions. Further, we present neutrophil-macrophage communications at the subset level and in specific ligand-receptor pairs, including IL6 to IL6 receptor signaling. Finally, we identified myeloid subset gene signatures in zebrafish that correspond to burn severity in human patients.

## Materials and Methods

### Zebrafish husbandry and wounding procedures

Adult and larval zebrafish are maintained following the animal protocol M005405-R02-A01 approved by the University of Wisconsin-Madison Institutional Animal Care and Use Committee (IACUC). Embryos were collected from adult *Tg(mpx:dendra,mpeg1.1:mCherry)* fish using natural breeding procedures and kept in E3 medium with 1% methylene blue inside a 28.5°C incubator. At 3 dpf, larvae were anesthetized with tricaine methanesulfonate (MS222, 200 mg/L) and wounded at the tail fin region using a handheld cauterizer. As the cauterizer wire heats up the E3 medium at close vicinity of the tail fin, the affected tissue area changes its transparency and shows as a wave spreading towards the body. The degree of burn wounding is controlled by turning off the cauterizer right before the wave of transparency change hits the tip of the notochord region. Post wounding, larvae were washed in E3 medium without MS222 and sent back to the incubator till the time of sample collection.

### Sample collection and library construction

At 6/24/48 hour-post-burn (hpb), burned and unwounded larvae at matching developmental stages (3/4/5 day-post-fertilization (dpf)) were transferred to 35 mm dish containing calcium-free PBS for 15 min and then anesthetized with tricaine methanesulfonate (MS222, 200 mg/L). A total of 150 fish were used for each time point by condition. Each dish of fish was digested with 2 mL digestion solution (0.25% trypsin, 1mM EDTA in PBS) at 28.5°C for 90 min with gentle pipetting every 10 min. Digestion was stopped by adding 200 mL digestion stop solution (1mM CaCl_2_, 100% FBS). Dissociated cells were filtered with 40 mm cell strainers and centrifuged for 3 min at 3,000 rpm at 4 °C. Cell pellets were resuspended in PBS with 10% FBS plus DAPI (1mg/mL).

Fluorescence activated cell sorting (FACS) was performed at University of Wisconsin Carbone Cancer Center Flow Lab using BD FACSAria. Targeted cells passed gating for cells versus debris, singles versus doublets, live versus dead and were high in either 488 nm/561 nm channel. Both dendra+ (488 nm channel) and mCherry+ (561 nm channel) cells were sorted into PBS with 10% FBS. Post sorting, cells were directly used for library construction in a Chromium controller at the Gene Expression Center of the University of Wisconsin-Madison Biotechnology Center (RRID: SCR_017757). Samples were processed with Chromium Single Cell Gene Expression Solution 3’ v2 (10X Genomics).

### Zebrafish scRNA-seq sequencing, data processing and quality control

Libraries were sequenced at DNA Sequencing Core of the University of Wisconsin-Madison Biotechnology Center (RRID: SCR_017759) on NovaSeq6000 system in S1 flow cells with read lengths of 29-bp + 90-bp (Read1 + Read2). Raw sequencing reads were processed by *cellranger count* in Cell Ranger (v6.1.2, 10x Genomics) using GRCz11 zebrafish genome with an improved gene annotation (13). Filtered cell-by-gene matrices generated were used for downstream analysis in R with Seurat (v4.3.0) (14). We filtered out cells with less than 200 genes detected or mitochondrial gene ratio greater than 20%. Doublets were removed using DoubletFinder with an estimated ratio of 2.3% for T1 samples (6hpb, 3dpf), 4.6% for 24 hpb, 6.1% for 4 dpf, 6.9% for 5 dpf, and 7.6% for 48 hpb (15).

### Unsupervised clustering and myeloid population selection

Individual Seurat objects were log-normalized and merged as one data object for linear dimensional reduction through obtaining top 2000 highly variable genes (*FindVariableFeatures*), data scaling and centering (*ScaleData*), and performing principal component analysis (*RunPCA*). To account for batch effects, the *Harmony* batch correction method was employed, using the individual sample label as the batch variable (16). Non-linear dimensional reduction was performed using Harmony-corrected top 100 PCs to generate UMAP axes (*RunUMAP*). Major cell types were identified by Louvain graph clustering at resolution of 0.1 on the nearest neighbors graph also constructed by corrected PCs (*FindNeighbors* and *FindClusters*). Neutrophils, macrophages, and cycling myeloid cells were annotated based on the top differentially expressed genes identified by Wilcoxon rank-sum test (*FindAllMarkers* with log2 fold change threshold of 0.25 and positive only) and subsetted for sub-clustering.

Isolated myeloid cells were split back into a list of Seurat objects by their original sample identity and normalized through *SCTransform*. The list was integrated based on the SCT assay and then through dimensional reduction by PCA and UMAP with top 30 PCs. Clustering was performed with top 30 PCs followed by differential expression analysis as described above.

### Human-zebrafish scRNA-seq label transfer

On the human data side, we received the processed Seurat object provided by the authors from Montaldo et al., and downsampled to keep 2654 cells in each neutrophil maturation stages (Precursor, Early Immature, Immature, and Mature) (6). On our zebrafish data side, we converted zebrafish gene symbols to their corresponding human orthologs by using the DIOPT scoring API with zebrafish as the source organism and human as the target organism (17). Only genes with a DIOPT score over six were kept ensuring confidence in ortholog calling. For genes with many-to-one orthologs, we used the most highly expressed gene from the zebrafish dataset. All label transfer procedures were conducted with tools in Seurat (v4.3.0). *SCTransform* was performed for both the received human neutrophil data and our own zebrafish neutrophil data. Anchors between the two dataset was identified by *FindTransferAnchors* using principal component analysis (PCA) with 30 PCs as reference reduction space, and SCT as normalization approach. Label transfer from zebrafish to human was performed by *TransferData* with the same dimensions used in anchor identification. Predicted zebrafish cluster name was added to human neutrophil dataset as a new metadata.

### ZebraHub and tessellated lymphoid network neutrophil visualization

Processed ZebraHub transcriptome data object covering all developmental time points (zf_atlas_full_v4_release.h5ad) was downloaded from https://zebrahub.ds.czbiohub.org/data and converted into H5Seurat format to be used in R environment with Seurat (v4.3.0). Only 5 dpf cells were used for visualizing general clustering and localization of myeloid populations on UMAP axes. Myeloid cells from all developmental time points were extracted by *CellSelector* based on myeloid marker expression pattern and used for co-expression visualization using *FeaturePlot* with blend = True.

Single-cell transcriptome data from adult zebrafish tessellated lymphoid network was processed as previously described (18). Neutrophil was annotated based on the top differentially expressed genes identified by Wilcoxon rank-sum test (*FindAllMarkers* with log2 fold change threshold of 0.25 and positive only) and subsetted for sub-clustering. Post neutrophil-only dimensional reduction, they were used for co-expression visualization using *FeaturePlot* with blend = True.

### Functional annotation for zebrafish myeloid subsets

We followed an analysis procedure adapted from Arora et al., with details listed below (19). Bulk level hierarchical clustering: For each cluster in neutrophils, we generated a pseudobulk profile by calculating the average expression level for each gene using the RNA assay. We performed *FindVariableFeatures* on neutrophils and filtered the average expression matrix to include only top 2000 highly variable genes. Expression levels of these genes were used for Pearson correlation calculation and hierarchical clustering.

Bulk level PCA: Pseudobulk profiles were generated similarly as above. We added one step before performing PCA to bring the average expression matrix back to log scale, which was missing in the original pipeline (19). These profiles were used as input for PCA with scale. = TRUE. Cell clusters were plotted onto two selected PC axes wrapped either by time point of sample collection to show burn-unwounded difference or by condition to show time course change. Top 500 genes ranked by positive or negative loading were extracted for GO enrichment analysis using Metascape v3.5.20230501 (20).

GSEGO: Gene ranking for each cluster was generated by conducting Wilcoxon rank sum test using *wilcoxauc* from presto v1.0.0 comparing between neutrophil subsets (21). By-cluster ranking was then taken as input for GSEGO analysis from clusterProfiler v4.6.2 with gene set sizes between 15 and 500 (22). Gene ontology terms were ranked by p-value and with the top five of them selected for visualization.

Expression of GO term associated genes: for each GO term of interest, all genes associated with it in each myeloid subset were pulled out. Average expression level of each gene in each subset by condition and time was normalized by the mean value from all cells and summed up to represent the expression level of corresponding GO term. These GO expression levels were converted to Z-scores for visualization.

### Pseudotime analysis

Cell trajectories were constructed using Slingshot v2.6.0 (23) or Monocle3 v1.3.1 (24). From neutrophil-only visualization, we found outliers not clustered with the rest of the neutrophils and removed them from trajectory analysis. UMAP embedding and neutrophil subset identities were used as input for Slingshot calculation.

For Monocle, we performed independent analysis from filtered matrices generated by *cellranger count*. Cell data set from each sample were combined into one data object for preprocessing, dimensional reduction, clustering, and graph identification using default parameters. Cells clustered with neutrophils labeled in the Seurat object were chosen for cell ordering with the root of trajectory manually selected at the Precursor-like_Neuts side.

### RNA velocity estimation

RNA velocity was calculated by velocyto v0.17.17 (25) using the default parameters from the alignment generated by *cellranger count* for each sample collected by condition and time. The loom files generated by velocyto were used as input in complement to the Seurat object containing only neutrophils for velocity estimation with scVelo v0.2.5 in Python (26). In brief, Seurat object and loom files were merged and filtered to contain only top 3000 genes with at least 20 counts shared between unspliced and spliced measurements. The resulting datasets were processed through *pca* and *neighbors* then for velocity estimation using the stochastic mode.

### Network analysis with CellOracle

The Seurat object containing all myeloid cells was converted to AnnData format then loaded in Python for network analysis with CellOracle v0.14.0 (27). The AnnData object was normalized by the total number of UMI counts per cell and filtered to include only the top 3000 genes with at least one count. Post filtering, the AnnData object was renormalized and used for *perform_PCA* followed by *knn_imputation*. For gene regulatory network (GRN) construction, we used the zebrafish promoter base GRN (danRer11_CisBPv2_fpr2) provided by the package. Cluster-specific GRN was constructed with top 2000 source-target links kept based on edge strength. All transcription factors (TFs) in the top 2000 links from all myeloid subsets were merged and clustered based on their degree of centrality. Top 30 TFs were included for visualization. The expression level of genes targeted by these TFs were extracted using the same approach as in **Functional annotation for zebrafish myeloid subsets**.

### Cell-cell interaction analysis with CellChat

Cell-cell interaction analysis was performed using the CellChat v1.6.1 R package (28). All myeloid populations were included for analysis. Functions from CellChat were used to identify changes in interactions between populations across conditions at comparable timepoints. Visualizations were constructed from built-in visualization functions in CellChat or through export of CellChat data and visualization with ggplot2.

### Scoring human burn microarray data with zebrafish gene candidates

Burn microarray data from SYSCOT study group was first stratified based on patient total burn surface area (TBSA) (29). Patients with 0-10% TBSA were classified as low TBSA, 11-20% were moderate TBSA, 21-30% were high TBSA, and 31%+ were very high TBSA. Using the human gene converted Seurat object constructed from **Human-zebrafish scRNA-seq label transfer**, we identified zebrafish myeloid subset markers in the form of their human orthologues with *FindAllMarkers* implementing MAST hurdle model. Differentially expressed genes were further filtered based on a log2 fold change of 0.5 and an adjusted p-value of 0.01.

Microarray datasets were then evaluated at a per-sample level by averaging the top 50 gene expression values from each subset specific signature. In the case where a subset gene signature contained fewer than 50 genes, all of the genes were used. The distribution of gene signature scores was visualized as raincloud plots with their means compared using a Wilcoxon rank-sum test as implemented in the ggpubr library function *stat_compare_means*. Null datasets were generated through 10000 permutation tests where TBSA labels were shuffled, and gene signature averages were calculated. The spearman correlation between individual genes within the found signatures and the subset gene expression signature averages were calculated with the base R *cor* function with the method parameter equal to spearman.

### Flow cytometry of human blood samples

Peripheral blood was collected from de-identified healthy donors and burn patients (Supplemental Table 1). Burn samples (blood) were collected within one week of hospital admission. All incubation and washing steps are performed at room temperature. Samples were incubated with red blood cell lysis buffer (BioLegend) diluted 1:10 in DI H2O for 10 minutes within 1 hour of sample collection and then washed twice with Cell Staining Buffer (BD Biosciences). For cell surface staining, Human TruStain FcX (BioLegend) was added at 1:20 to block Fc receptors for 10 minutes, followed by incubation in Zombie NIR Live/Dead stain at 1:1000 (BioLegend) for 10 minutes. Protein-targeted staining was done with the following antibodies in Cell Staining Buffer: Alexa Fluor 700-conjugated anti-human CD16 at 1:400, PE/Fire-640-conjugated anti-human CD66b at 1:400, PerCP-conjugated anti-human CD3 at 1:200, PerCP-conjugated anti-human CD19 at 1:200, PerCP-conjugated anti-human CD56 at 1:400, PerCP-conjugated anti-human CD203c at 1:100 (BioLegend), BUV395-conjugated mouse anti-human CD45 at 1:100 (BD Biosciences), and PE-conjugated GPR84 at 1:200 (Alomone Labs). After staining for 30 minutes, cells were washed with Cell Staining Buffer and resuspended in Fixation Buffer (BioLegend) for 15 minutes. Post fixation, cells were washed and resuspended in 300 uL Cell Staining Buffer and filtered through a 70-micron filter. Flow cytometry data was collected on the Cytek Aurora flow cytometer. FCS files were exported and analyzed using FlowJo software v.10.9.0. Neutrophils were identified as CD3/CD19/CD56/CD203c-, CD45/CD66b/CD16+ cells with granulocyte scatter. These cells were then evaluated for the expression of GPR84. Fluorescence minus one (FMO) of GPR84-PE was used as control to quantify cells expressing GPR84.

## Results

### Single-cell transcriptomics reveals myeloid cell heterogeneity in larval zebrafish

To characterize the molecular heterogeneity of myeloid cells in larval zebrafish, we used transgenic zebrafish *Tg(mpx:dendra,mpeg1.1:mCherry)* with fluorescently labeled neutrophils and macrophages. Cells were collected from both burned and unwounded larval zebrafish along a time course of three days (Figure 1A, Materials and Methods). We chose three time points to capture burn wound healing (6 hpb, 24 hpb, 48 hpb. hpb: hours post burn) that covers the myeloid cell recruitment and inflammation resolution stages (10). In addition, we included the corresponding time points in normal development as unwounded controls (3 dpf, 4 dpf, 5 dpf. dpf: day post fertilization). We obtained transcriptomes of all cells at single cell resolution using a droplet-based 3’ mRNA capturing approach from 10X Genomics. Initial processing revealed six known major cell populations, including neutrophils (*mpx, lyz*), macrophages (*mpeg1.1, marco*), lateral line cells (*prox1a, f11r.1*), epithelial cells (*cldnb*, *krt4*), erythrocytes (*hbae1.1, hemgn*), photoreceptors (*opn1mw1, gnb3b*), and one cycling myeloid population with both neutrophil and macrophage signatures and markers labeling cell cycle (*mki67*, *top2a*). One identified population has not been previously characterized, with higher expression of *cebpd* and *her6* (Supp. Figure 1A-B). We focused on neutrophils and macrophages (a total of 19118 cells) for further clustering analysis that identified six macrophage subtypes and four neutrophil subtypes (Figure 1B, Supp. Figure 1C, Supp. Figure 2). Two of the six macrophage subsets showed enriched expression of cell cycle related genes, including Cycling_Mac1, which had high level scoring for both S phase and G2M phase markers (Supp. Figure 3).

**Figure 1.**
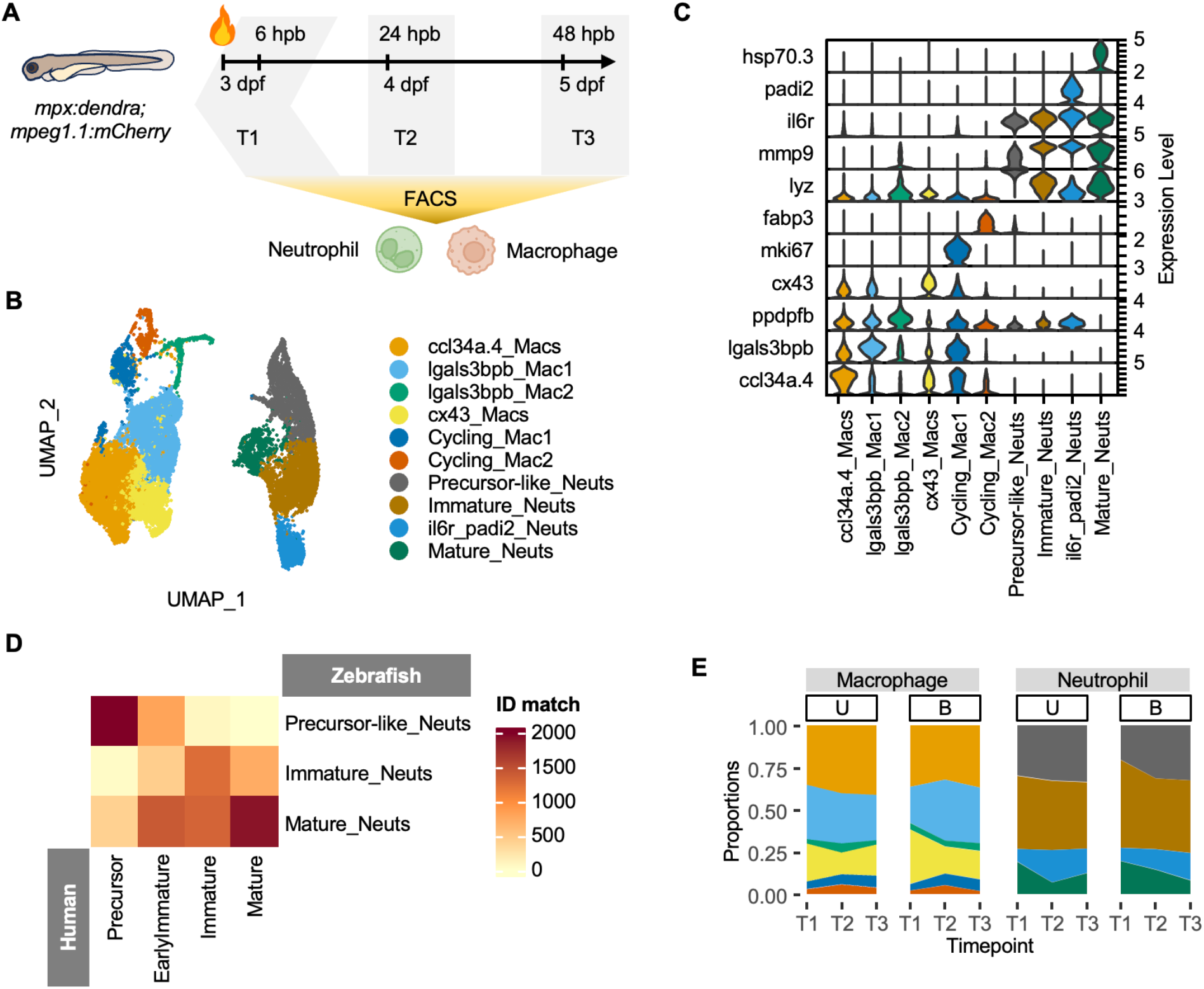
Myeloid population structure in larval zebrafish under homeostasis and burn wounding. (A) Overall experimental design. Three timepoints were collected for either burn (6/24/48 hpb) or unwounded (3/4/5 dpf) conditions from larval Tg(*mpx:dendra*;*mpeg1.1:mCherry*). Neutrophils and macrophages were enriched from Fluorescence-activated Cell Sorting (FACS). Collected cells were used for single cell RNA-seq library generation and sequencing. (B) Myeloid cell subsets visualized on UMAP axes. In total, there are four neutrophil subsets and six macrophage subsets. (C) Distribution of representative marker expression across subsets shown as violin plot. (D) Identity matching between human neutrophil stages and zebrafish neutrophil clusters post label transferring. (E) Proportion of each myeloid subset by condition and time points. U: Unwounded, B: Burn.

Current molecular annotation for myeloid subsets in zebrafish is still limited, although there has been a recent report from adult zebrafish bone marrow (12). Therefore, we named clusters both by known annotations and their representative top expressed genes. For macrophage subsets (Figure 1C), we examined the expression distribution of known M1/M2 subtype markers and could not match zebrafish subsets with this polarization nomenclature (Supp. Figure 1D). Based on gene signatures, we named the macrophage clusters as ccl34a.4_Macs, lgals3bpb_Mac1, lgals3bpb_Mac2, and cx43_Macs (Figure 1B, Supp. Figure 1C, Supp. Figure 2). Among which, *ccl34a.4*+ and *lgals3bpb*+ macrophages were also defined in a recent macrophage scRNA-Seq datasets from adult zebrafish under homeostasis (30). For neutrophils, we performed cross-species label transferring between our zebrafish data and a comprehensive scRNA-Seq mapping of human neutrophils, including four stages: precursors, early immature, immature, and mature neutrophils (6). We mapped zebrafish neutrophil subsets to human neutrophil stages using homologs of marker genes (Methods & Supp. Figure 4A-B) and identified three out of four zebrafish neutrophil subsets that align well with human neutrophil stages; named as Precursor-like_Neuts, Immature_Neuts, and Mature_Neuts respectively (Figure 1D, Supp. Figure 4C). The *il6r* and *padi2* expressing neutrophil subset could not be mapped to any of the human neutrophil states, thus keeping its marker-based name as il6r_padi2_Neuts. We confirmed the existence of this population in scRNA-Seq data from the ZebraHub database (Supp. Figure 5 A-C), which covers up to 5 dpf, and in the adult zebrafish tessellated lymphoid network (Supp. Figure 5 D-E) (18).

All subsets exist in both burned and unwounded conditions at each time point (Figure 1E). This consistent distribution suggests that as early as 3 dpf, larval zebrafish have established heterogeneous populations of myeloid cells that sustain through 5 dpf. In addition, burn wounding does not eliminate or induce any subset. In the burn healing process, Precursor-like_Neuts and lgals3bpb_Mac1 showed more apparent increases at an early time point compared to unwounded control (Supp. Figure 1E). On the contrary, Mature_Neuts and cx43_Macs showed a more apparent decrease in burn. Taken together, we demonstrate the existence of heterogeneity within myeloid populations in both normal development and during burn wound healing in larval zebrafish. Hereafter, we focus more on dissecting the molecular features of neutrophil subsets and their interactions with macrophages.

### Neutrophils present functional diversity with temporal and conditional specificities

We next aimed to understand the function of neutrophil subsets in normal development and burn wounds. First, to identify the global differences across subsets, conditions, and time points, we summarized expression in each subset and performed hierarchical clustering among subsets (Figure 2A). Precursor-like_Neuts are grouped together, regardless of their condition or time point of collection. However, Immature_Neuts and il6r_padi2_Neuts at the earliest time point, either in burn or unwounded condition, are grouped with the Mature_Neuts subset. To further dissect the variance between subsets, we performed principal component analysis (PCA) on the pseudobulk profiles. The top two PCs explained 24% of variance (Figure 2B). By examining the grouping of subsets based on the first six PCs, we found that PC2 correlates with time progression and separated burned from unwounded at T1, while PC5 better separated different subsets (Supp. Figure 6).

**Figure 2.**
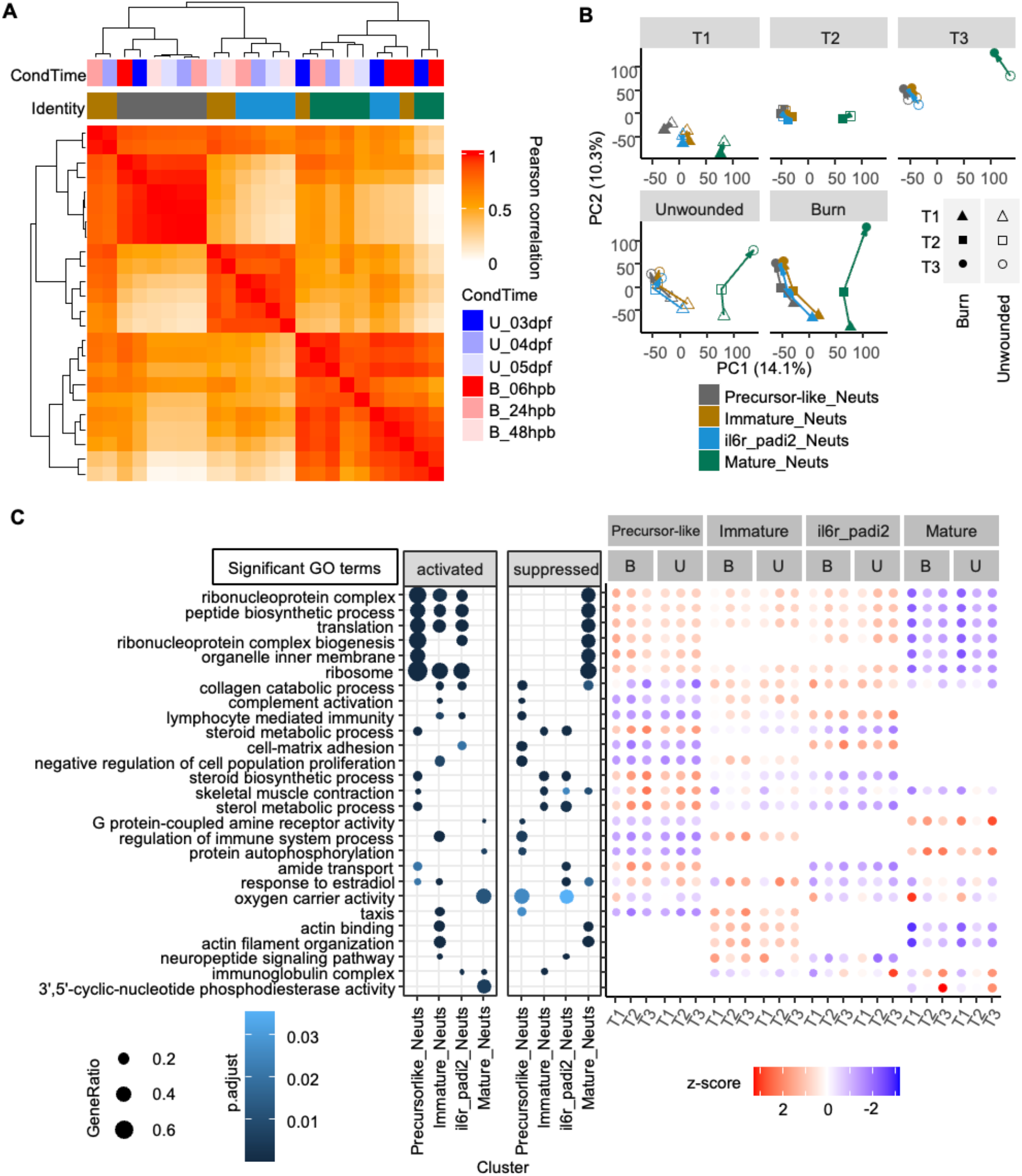
Neutrophil subset functional annotations. (A) Hierarchical clustering of neutrophil subsets. Neutrophil subsets are grouped by conditions and time points to create pseudobulks for Pearson correlation calculation. Identity color coding is the same as in (B). (B) Neutrophil subset clustering pattern in Principal Component Analysis (PCA). Pseudobulks are shown on PC1 and PC2 axes with the upper panel grouped by time points and the lower panel grouped by conditions. Arrows in the upper panel point to the Burn condition, while those in the lower panel follow the time flow. (C) GO term enrichment in neutrophil subsets and the expression level of corresponding genes. Gene ratio and adjusted p-value of each GO term listed on the left are shown as size and color range in the left hand side dot plot, with the activated and the suppressed listed separately. Z-score of GO-associated genes are shown as color in the right hand side dot plot, with neutrophils grouped by subsets, condition, and time.

Next, we asked what drives the separation on PC axes. We performed Gene Ontology (GO) enrichment analysis on the top 500 genes with positive or negative correlation with each PC loading for PC annotation (20) (Supp. Figure 7). Genes with negative loading of PC2 showed enrichment for both active protein synthesis (dre04141 Protein processing in endoplasmic reticulum, R-DRE-532668 N-glycan trimming) and degradation functions(GO:0030163 protein catabolic process, GO:0006517 protein deglycosylation). This suggests a higher turnover rate for neutrophils in T1 (3 dpf/6 hpb) than later time points and in burn versus unwounded condition at T1. Genes associated with PC5 enrich for terms that reflect differences between subsets, with the positive direction associated with the Precursor-like_Neuts, showing cell cycle features (GO:0006260 DNA replication, dre03410 Base excision repair, R-DRE-69206 G1/S Transition), while the negative direction showed more mature cell functions (GO:0045321 leukocyte activation, GO:0050900 leukocyte migration) and the signaling pathways that mediate them (GO:0007229: integrin-mediated signaling pathways, GO:0034097 response to cytokine, GO:0007179: TGF beta receptor signaling pathways). The data suggest that from Precursor-like_Neuts, to Immature_Neuts, then to il6r_padi2_Neuts, the cell cycle-related signature drops while immune activity increases.

While PCA provides a global separation between subsets, it is not enough for comprehensive functional annotation for each. To increase the granularity of functional annotation, we performed gene set enrichment analysis (GSEGO) for all subsets (Figure 2C) (22). Compared to over-representation in GO enrichment analysis, this approach allowed for both the existence and ranking of genes to be taken into consideration. Zebrafish GO terms were used as gene sets for rank testing. We further complemented the enrichment analysis with the average expression level of the genes associated with each GO term in each subset at individual time points (Materials and Methods). We found subset-specific enrichment in certain biological processes, including translation-related terms in Precursor-like_Neuts, Immature_Neuts, and il6r_padi2_Neuts (GO:1990904 ribonucleoprotein complex, GO:0043043 peptide biosynthetic process). Precursor-like neutrophils are also highly enriched in steroid metabolism-related biological processes (GO:0008202, GO:0006694, GO:0016125), but without significant expression level changes between burned and control conditions. Both Immature_Neuts and il6r_padi2_Neuts enrich for collagen catabolic process (GO:0030574). Genes associated with this term are also highly expressed in them, with highest levels at 6 hpb condition (T1 in burn condition). This suggests potential early involvements of Immature_Neuts and il6r_padi2_Neuts in the wound healing processes. Immature_Neuts also uniquely enriched for neuropeptide signaling (GO:0007218), suggesting that zebrafish might have a subset of neutrophils that are regulated by neuropeptides like humans (31). Mature_Neuts enriched for GPCR activity (GO:0008227), protein autophosphorylation (GO:0046777), oxygen carrier activity (GO:0005344), immunoglobulin complex (GO:0019814), and cyclic nucleotide degradation (GO:0004114). Among these terms, GPCR activity and cyclin nucleotide degradation activity has their associated genes more highly expressed in 48 hpb condition (T3 in burn). Cyclic adenosine monophosphate (cAMP), as one type of cyclic nucleotides, is known to be at high levels in the urine of severe burn patients that gradually drops to a level closer to that in mild burn patients (32). It acts downstream of prostaglandin E2 (PGE2) to suppress the activation and growth of T-lymphocytes and granulocyte-macrophage progenitors (33, 34). Mature_Neuts in zebrafish could be involved in the attenuation of the cAMP-mediated immunosuppression effect at a later stage in burn wound healing. These results indicate that zebrafish neutrophil subsets play distinctive roles during normal development and could selectively enhance specific functions during burn wound healing.

### Neutrophil state transition in zebrafish is conserved with humans

From zebrafish-human label transfer (Figure 1), we learned that zebrafish neutrophils follow the same path as human neutrophils through maturation. But we also saw that il6r_padi2_Neuts do not match with human neutrophil states. What’s the relationship between this unique subset and the conserved differentiation path? We first examined the state transition paths through two independent approaches: trajectory reconstruction with pseudo-time analysis (slingshot, monocle) and RNA velocity estimation (Materials and Methods). Both slingshot and monocle inferred trajectories consistently predict a branched trajectory, with one branch matching the states of human neutrophils developing from Precursor-like_Neuts to Immature_Neuts then to Mature_Neuts (Figure 3A, Supp. Figure 8A). Il6r_padi2_Neuts are on the other branch of the trajectory that connects with Immature_Neuts, but with unclear directionalities. We examined the expression of *mmp9*, a marker associated with zebrafish neutrophil maturity, and found that il6r_padi2_Neuts have a higher level than Mature_Neuts (Supp. Figure 8B) (12). However, human homologs of the top-ranking signatures of il6r_padi2_Neuts are mainly expressed at the early immature or immature stage in human neutrophils (Supp. Figure 8C).

**Figure 3.**
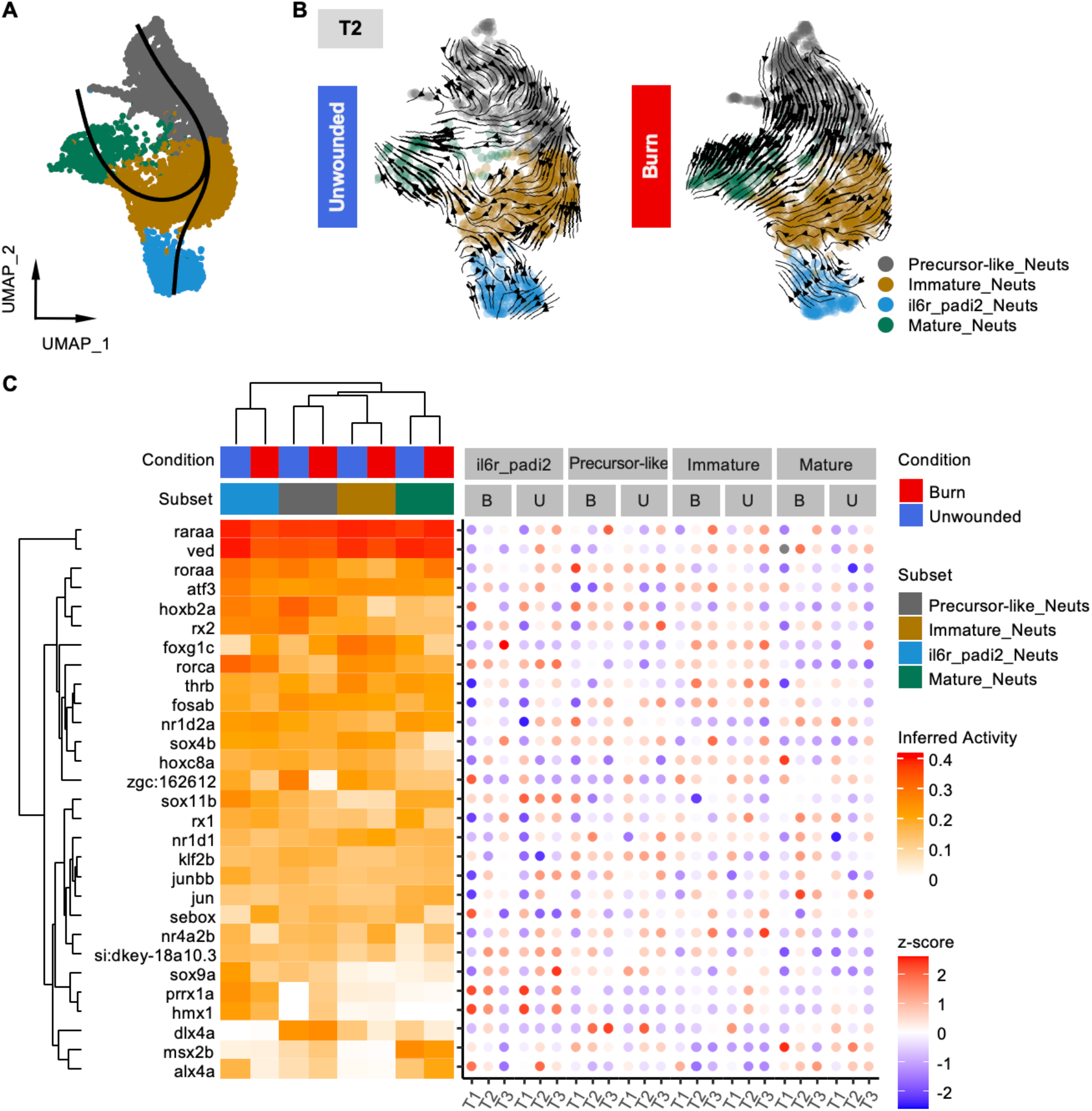
State transition between neutrophil subsets. (A) Slingshot predicted trajectory on the same UMAP axes as in Figure 1B, with neutrophil only. (B) Velocity prediction for neutrophils at T2 (unwounded: 4 dpf, Burn: 24 hpb) projected as arrows on the same UMAP axes as in (A). (C) Degree of centrality of top 30 transcription factors neutrophil subsets (left) complemented with the expression level of their downstream genes (right). Level of degree of centrality marked as Inferred Activity.

To further understand the direction of the transitions among subsets, we performed RNA velocity analysis in neutrophils (Figure 3B, Supp. Figure 8D) (Materials and Methods). RNA velocity incorporates the dynamics of the spliced and unspliced forms of mRNA across genes to infer the future state of a cell (25, 26). While the differentiation path matches the predicted directionality and remains stable, the path connecting il6r_padi2_Neuts with Immature_Neuts could be lost. This suggests that the il6r_padi2_Neuts might not be the terminal state of the path like the Mature_Neuts. Comparing between conditions, we found a strong velocity trend from Precursor_Neuts to Immature_Neuts and Mature_Neuts in burn, especially at T2 (Figure 3B). This suggests a potential trauma-induced emergency hematopoiesis in zebrafish in response to burn wounding. Connecting this to the proportional changes shown in Figure 1E, we see the transition resulted in expansion of Precursor-like_Neuts and il6r_padi2_Neuts and a shrinkage of Mature_Neuts.

We next aimed to identify what factors drive the state transition of neutrophil subsets. We performed network analysis using CellOracle, which takes advantage of both open chromatin signature and gene expression at single-cell resolution to build cell-state-specific Gene Regulatory Networks (GRNs) (27). We utilized the built-in promoter base GRN models in CellOracle to compensate for our lack of matching chromatin accessibility profiles. We constructed a transcription factor (TF) to target gene link and compared the degree centrality for TFs in different subsets across conditions and time points (Supp. Figure 9A, Figure 3C). The degree centrality level reflects the number of downstream targets a TF might control and is used here to infer TF activity. Macrophages were included for elucidating cell-type specific TF activity or excluded to focus on the between-subset comparisons in neutrophils. We found *raraa* and *ved* are more active in neutrophils, while *foxg1c*, *sox9b*, and *rxraa* are in macrophages (Supp. Figure 9A). As expected, between-subset distinctions are less pronounced compared to that from between cell types. To further understand neutrophil subset-specific TFs across conditions and time, we calculated the average expression level of predicted downstream genes in neutrophil subsets for each candidate TF (Figure 3C). In general, TFs with higher inferred activity in a subset also presented a relatively higher expression level of downstream target genes in that subset. This suggests that active TFs are not only controlling more genes, but also enhancing their expression level. Mature_Neuts and il6r_padi2_Neuts are the two branches in the trajectory analysis, so we compared between the two and identified TFs with their activities associated with these populations. We found TFs associated with one branch independent of wounding conditions (*msx2b* vs *prrx1a*, *hmx1*, *hoxb2a*, and *rx2*), and TFs that showed condition-dependent patterns (Supp. Figure 9B). Most of the candidates do not have known neutrophil-associated functions, but are associated with retinoic acid (RA) signaling, including suppression (*alx4a*, *dlx4a*, and *sox9a/b*) (35–37), activation (*hmx1*, *sox4b*, *msx2b*, *hoxb2a*, *nr6a1a*) (38–42) and serving as co-receptors (*nr4a2b*) (43). Altogether, we identified the state transition paths in zebrafish neutrophils compared to human neutrophils and identified TFs that potentially drive the deviations.

### Burn injury induces neutrophil-macrophage communication

Cells in a living organism communicate with one another to adjust their function to their cellular microenvironment. In search of cell-cell communication pathways of early innate immune responses that are activated or suppressed in burn wounding conditions, we performed CellChat analysis using all myeloid subsets (28). We converted a curated human ligand-receptor pair database for use in zebrafish to infer outgoing and incoming signals in each population in a context-dependent manner as described in the methods. At the global level, the total number of interactions between myeloid subsets in both burned and unwounded conditions first drops, then increases as time goes from T1 to T3 (Figure 4A-B). However, the average strength of these interactions, measured by co-expression of Ligand and Receptor genes, presents a condition-specific pattern, where it mostly remains high in burned samples, but rapidly decreases in the unwounded condition. Zooming in on the subset level, we found a differential contribution from myeloid subsets to the overall level of outgoing and incoming signals in the burned versus unwounded comparison (Figure 4C). At T1, Precursor_Neuts, Immature_Neuts, and Mature_Neuts follow the same enrichment pattern, with relatively higher signals in both outgoing and incoming directions in burn. At T2, all neutrophil subsets picked up this pattern. In addition, all subsets except for lgals3bpb_Mac2, showed relatively higher incoming signals in burn conditions. At T3, all myeloid subsets except for Mature_Neuts and lgals3bpb_Mac2 showed incoming enrichment in the burn condition, while all except for Cycling_Mac1 and lgals3bpb_Mac1 showed outgoing enrichment in the burn condition. Overall, myeloid cells in burn had extended signaling interactions until 48 hpb, with neutrophils showing more striking differences between conditions.

**Figure 4.**
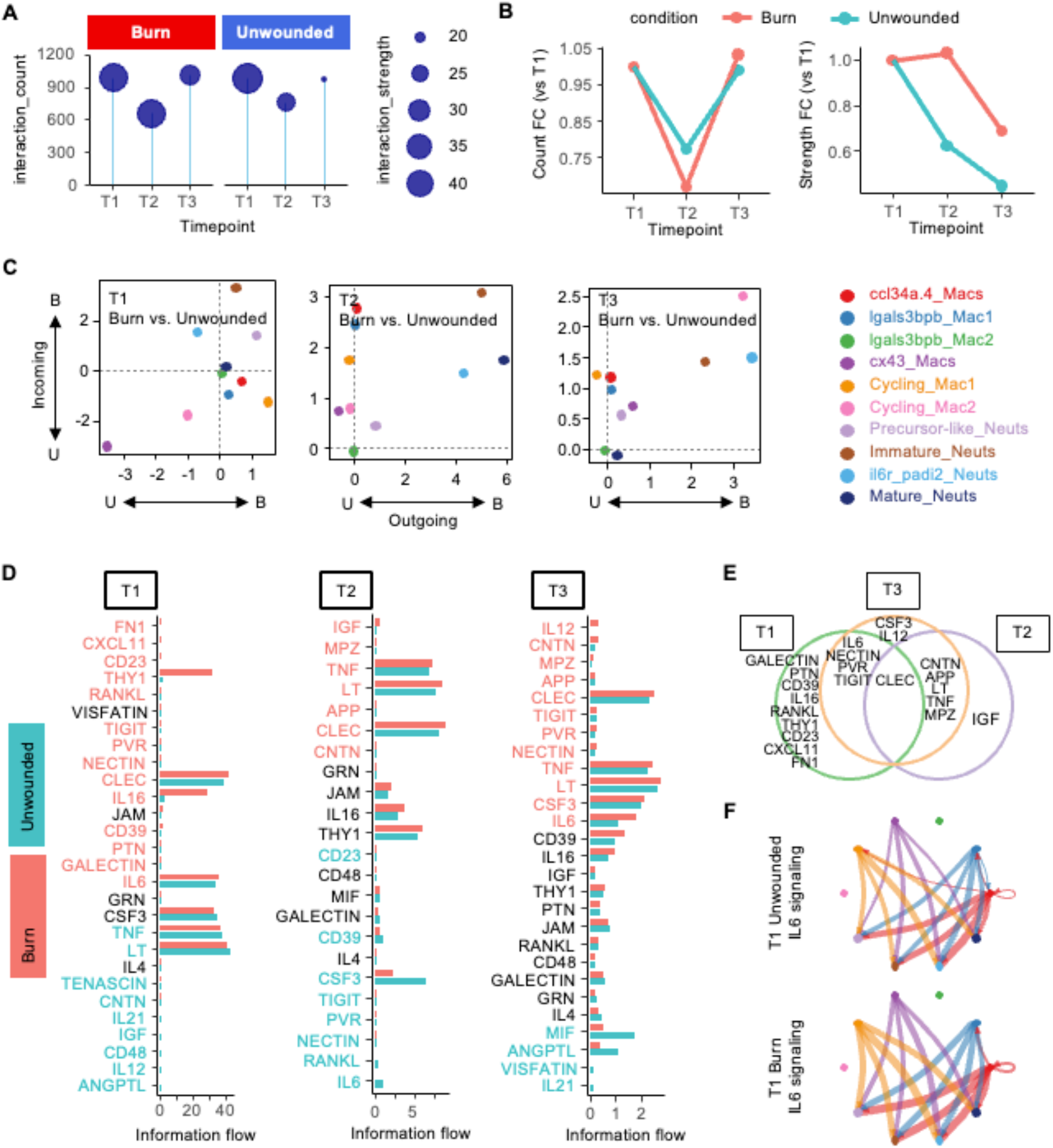
Cell-cell communication across myeloid subsets. (A) Interaction measurement between myeloid cells by condition and time. Total number of interactions are shown on the Y-axis, while interaction strengths are shown by the size of circles. (B) Relative interaction counts and strength compared to T1 calculated using data from (A). (C) Overall signaling level by population at each time point. X-axis measures the outgoing signals, Y-axis measures the incoming signals. The direction of the arrows show the direction of enrichment. (D) Signaling information flow by ligand-receptor pairs ranked by the burn-to-unwounded ratio and the significance level. The labels of the ligands are color-coded by their bias towards one of the two conditions. (E) Venn plot presenting common and unique candidate ligand-receptor pairs across time points. (F) Myeloid subset signaling network of IL6 pathway. Population color coding is shared with (C).

We next determined the key ligand-receptor pairs that contribute to the burn-induced interactions. We calculated information flow and ranked each ligand-receptor pair by the differences between burn and unwounded conditions and the statistical significance of the comparison (Figure 4D). We looked for common and unique ligand-receptor pairs that are relatively enriched in burn conditions across time points (Figure 4E). Then, we narrowed down our search for candidate pathways by focusing only on the ones with high levels of overall information flow. Among the candidate pathways, *THY1* and *IL16* signaling both showed a high burn-to-unwounded ratio only at early time points, with the former in the incoming direction, while the latter in the outgoing direction for neutrophils (Figure 4D, Supp. Figure 10A). Ligands of *THY1* bind integrin and play a role in mediating endothelial adhesion and migration of myeloid cells (44–46). Binding of recombinant human *THY1* induces *MMP9* expression in neutrophils, which in turn enhances their migration ability (47). This aligns with the collagen metabolic function enrichment pattern in Immature_Neuts and il6r_padi2_Neuts, suggesting that their migration regulation is conserved with humans and mice. The *IL6* pathway presents a trend-switching pattern, where it shows higher information flow in burn conditions in T1 and T3, but lower in T2. By examining the networks formed by *IL6-IL6R* interactions among myeloid subsets, we found that the *IL6* signal is mostly sent out by macrophages and received by neutrophils (Figure 4F, Supp. Figure 10B). Previous work using zebrafish *il6r* mutant has shown that the *il6-il6r* axis is essential for neutrophil but not macrophage recruitment to the burn wound area (11). This matches with the high-in-burn pattern for *IL6-IL6R* information flow at T1 and T3. T2 (24 hpb) is the time point when the majority of the neutrophils would have migrated out of the wound site in wild type larval zebrafish post burn wounding. Therefore, we think that the reverse pattern in T2 might suggest the need for *il6-il6r* signal attenuation for inflammation resolution. This finding points to the hidden complexity of the signaling pathways at myeloid subset level.

### Zebrafish myeloid subset signatures associate with human burn severity

To test if our findings in zebrafish burn models could translate to human burns, we examined the association between human orthologs of zebrafish myeloid subset signatures and burn severity in human patients using published microarray data (Figure 5A) (29). We extracted top expressed genes from myeloid subsets in zebrafish and converted those to human orthologs using the DIOPT score (17). To gain a refined range of human burn severity measured by Total Body Surface Area (TBSA), we divided the TBSA levels to four categories: 0-10% (Low), 11-20% (Medium), 21-30% (High) and 31-40% (Very High) with >100 samples per each category. First, we asked if any myeloid subset signatures are associated with the TBSA levels and found that Immature_Neuts signatures positively associate with TBSA, while lgalsbpb_Mac1 negatively associates with TBSA (p < 2.22e-16 for both subset comparing between Low and Very High TBSA, Figure 5B). Intriguingly, the trend is preserved with only samples collected at day 1 (with sample size reduced by half. Both p < 1e-3 comparing between Low and Very High TBSA), indicating this might be a signature of early innate response in human burns. To ensure the correlation was not observed due to the large sample size, we permuted TBSA levels for 10,000 times and found no correlation (p>0.1) as expected. Furthermore, we aimed to identify if there are any individual genes that correlate with the TBSA levels and found *GYG1*, *GPR84*, *CAPG*, and *MMP8* positively correlated with TBSA levels (Figure 5C), while *CD74*, *RUNX3*, and *PSAP* negatively correlated with TBSA levels among the top 50 genes in each subset.

**Figure 5.**
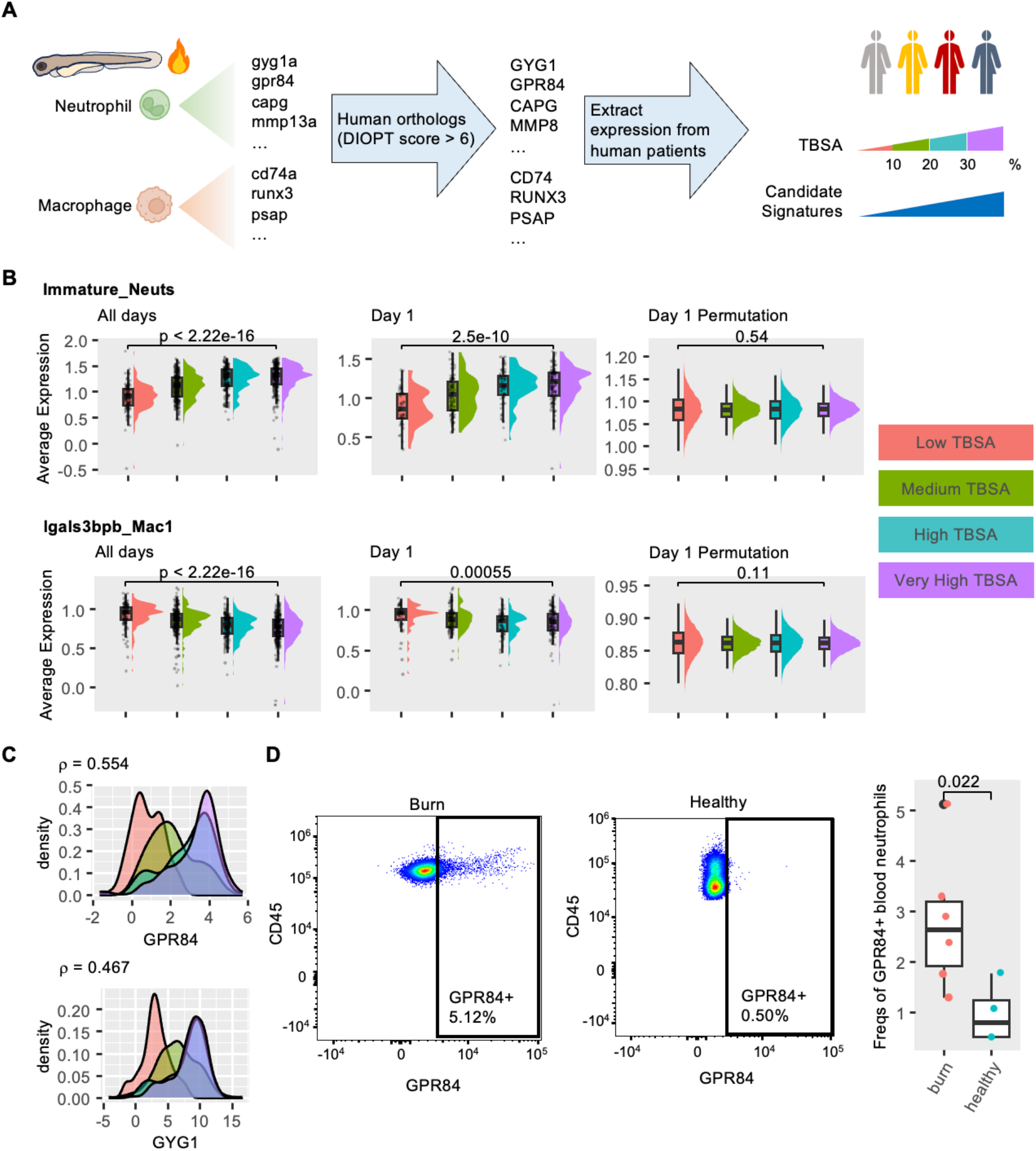
Zebrafish myeloid signatures in human burn patients. (A) Schematic view of validating signatures identified from zebrafish myeloid subsets in human burn patient expression profile. (B) Expression level distribution of Immature_Neuts signatures (top panel) and lgals3bpb_Mac1 (bottom panel) by patient TBSA shown as raincloud plots. Paired T-tests were performed between the Low and the Very High TBSA groups. (C) Expression distribution of GPR84 and GYG1 in patients by TBSA level. Spearman correlation (ρ) is calculated between gene expression and TBSA level. (D) Representative flow plots (left) and quantifications of frequencies shown as percentages (right) of GPR84+ neutrophils in burn (n = 6) and healthy (n = 4) blood. Paired samples t-test, p = 0.022.

Among these burn-severity-associated marker genes, GPR84, CD74, and PSAP all locate on the plasma membrane, making them directly detectable through flow cytometry. We used GPR84, a G_αi_-coupled receptor, with its activation by medium-chain fatty acids induces pro-inflammatory response in neutrophils (48), as the top candidate to test whether their RNA level in whole blood association with burn severity is reflected by protein level on the cell surface. We performed flow cytometry using peripheral blood collected from both burn patients and healthy donors with generally matching demographics (Methods; Supplemental Table 1). We found there is a higher frequency of GPR84+ neutrophils in the blood neutrophils of burn patients compared to that of healthy controls (Figure 5D, paired samples t-test, p = 0.022), suggesting its involvement in the human burn wound healing process. Together, this demonstrates the potential of translating zebrafish burn wounding research to human patient treatment.

## Discussion

Neutrophil heterogeneity at both the phenotypic and molecular level has been suggested by recent studies. However, to what degree such heterogeneity exists in tissue damage, in particular, in burn injury, has not been well characterized. In this work, we focused on the myeloid diversity in burn wound healing response using larval zebrafish as a model. By performing transcriptional profiling at single cell resolution, we identified four neutrophil subsets and six macrophage subsets in both normal development from 3-5 dpf and burn wound healing from 6-48 hpb. These subsets are present at the earliest time point we captured in development, suggesting that fish have developed this diverse group of myeloid cells as early as 3 dpf. In addition, general population structure remains the same in burn wound healing compared to unwounded control. This consistent distribution of myeloid subsets suggests that burn wounds do not induce the generation of new myeloid populations.

Due to the limited molecular definition of neutrophils in zebrafish, we took advantage of published human and mouse transcriptomes for subset identity confirmation. Through label transfer between human and fish neutrophils, we found three out of the four neutrophil subsets in fish could map to human neutrophil stages. Among these, Mature_Neuts shows more identity overlap with human neutrophils not in the mature stage, suggesting that the zebrafish neutrophils could be less mature than the human counterparts. Trajectory analysis further confirms that these three subsets follow a maturation path similar to that in humans. This subset-level conservation makes fish-based wounding models for studying innate immunology more relevant to humans.

Il6r_padi2_Neuts, the neutrophil subset that does not score well in human neutrophil stages, lies on a branch separated from the trajectory of maturation. RNA velocity analysis also suggests that this population might not be an end state as we found with the Mature_Neuts. To further understand the factors driving the separation between il6r_padi2_Neuts and Mature_Neuts, we constructed gene regulatory networks for each and identified key TFs associated respectively. Most of the candidates or their homologs are known to be involved in retinoic acid (RA) signaling. Retinoic acid is essential for neutrophil maturation, as it has been studied in the context of acute promyelocytic leukemia (APL) patients, cell lines and knock-out mouse models (49). The subset-dependent RA signatures suggest that RA signaling might be involved in finer control of neutrophil maturation in zebrafish. Further work is needed to explore the il6r_padi2_Neuts and its development in zebrafish.

To understand the roles of these neutrophil subsets in burn wound healing, we performed functional annotation and compared between burned and unwounded conditions. Although general expression of GO term associated genes present a similar subset enrichment pattern in both conditions, time point-dependent burn response could be observed in certain terms. For instance, genes associated with collagen catabolic processes and cyclic nucleotide degradation are expressed at much higher levels in burned condition at 6 hpb and 48 hpb respectively. This suggests that burn could induce time-sensitive functional enhancement in neutrophil subsets.

Burn wounding not only affects functions of individual neutrophil subsets, but also changes the way they communicate. Cell-cell interaction analysis revealed more outgoing and incoming signaling from neutrophils in burned than in unwounded conditions. In particular, *THY1*, *IL16* and *IL6* stood out as candidate ligand-receptor pairs that possess higher information flow in the burned condition. Their enrichment patterns aligned well with functional study from humans and mice suggesting their roles in myeloid recruitment and migration. In our work, we relied on the co-expression patterns in myeloid subsets to infer the communication between them. However, other cell types could express the same ligands or receptors and be involved in the communication as well. Although other cell types were captured in the cell sorting, we chose to leave them out due to the potential bias in our sorting strategy to more likely be obtaining cells with auto-fluorescence from populations other than the labeled myeloid cells. To better characterize cell-cell interaction between neutrophil subsets and other cell types in burn wound healing, an all-inclusive profiling that samples other cell types in appropriate ratios is needed in future work.

As an end goal of this study, we evaluated the translational value of the larval zebrafish model for human burn wounding. We confirmed associations between zebrafish neutrophil subset signatures and human burn severity using a human microarray dataset. Although other factors such as presence of inhalation injury, comorbidities of the patient, and depth of burn also contribute, total burn surface area (TBSA) is still the major factor in determining severity of burn. These findings suggest the potential of using zebrafish as a model to study human burn responses at the molecular level. In fact, neutrophils carrying these signatures could be found in a much higher level in the peripheral blood of burn patients than healthy donors. With more in-depth characterization aligning burn wound healing time courses between humans and zebrafish, the zebrafish model will become more helpful in facilitating our understanding of cell type-specific responses in burn in humans.

## Acknowledgements

We thank all members of the Huttenlocher lab, the Dinh lab, and the Gibson lab for discussion and support in completing this work. We thank the Gene Expression Center at the University of Wisconsin-Madison for single-cell library construction and sequencing assistance, and the University of Wisconsin Carbone Cancer Center Flow Cytometry Laboratory, supported by P30 CA014520, for use of its facilities and services. The authors thank the University of Wisconsin Carbone Cancer Center BioBank, supported by P30 CA014520, for use of its facilities and services. We thank Dr. Montaldo Elisa and Dr. Ostuni Renato for providing us with processed data for comparing between human and zebrafish neutrophils.

## Author contributions

Y.H., A.H., and H.D. conceived the idea and designed the experiments for this project. J.R. and Z.S. performed the experiments. Y.H., P.K., A.G, Z.C. and H.D. analyzed and interpreted the sequencing data. Z.S, A.LF.G, Y.H., and H.D analyzed flow cytometry data. Y.H., P.K., and H.D. wrote the manuscript with feedback from all other authors.

## Data and materials availability

All raw and processed sequencing data generated in this study have been submitted to the NCBI Gene Expression Omnibus (GEO; http://ncbi.nlm.nih.gov/geo/) and will be available once the manuscript has been certified by peer review. Source codes and processed data will be provided at https://github.com/huydinhlab/HouKhatriZebrafishBurn

## Footnotes

1 **Funding**: Y.H. was supported through The American Association of Immunologists Intersect Fellowship Program for Computational Scientists and Immunologists. P.K. was supported by an NLM training grant to the Computation and Informatics in Biology and Medicine Training Program (NLM5T15LM007359). Research reported in this publication was supported by the National Institute Of General Medical Sciences of the National Institutes of Health under Award Numbers R35GM150893 (HD), R35GM118027 (AH). The content is solely the responsibility of the authors and does not necessarily represent the official views of the National Institutes of Health.

