## Supplemental materials for "Single-cell transcriptional landscape of temporal neutrophil response to burn wound in larval zebrafish"

**List of Supplemental Materials**

**Supplemental Figure 1 Myeloid cell identity confirmation.**

**Supplemental Figure 2 Representative expression distribution of myeloid subset signatures on UMAP axes.**

**Supplemental Figure 3 Cell cycle features in myeloid subsets.**

**Supplemental Figure 4 Identity label transfer between human neutrophil stages and zebrafish neutrophil subsets.**

**Supplemental Figure 5 Confirm il6r_padi2_Neuts in ZebraHub and tessellated lymphoid network.**

**Supplemental Figure 6 Neutrophil subsets on PC axes.**

**Supplemental Figure 7 Functional annotation of PC associated genes.**

**Supplemental Figure 8 Orthogonal trajectory analysis of neutrophil subsets.**

**Supplemental Figure 9 Transcription factor comparison among myeloid subsets.**

**Supplemental Figure 10 Pathway-specific network.**

**Supplemental Table 1 Patient metadata for blood sample collection**

Supp. Fig. 1

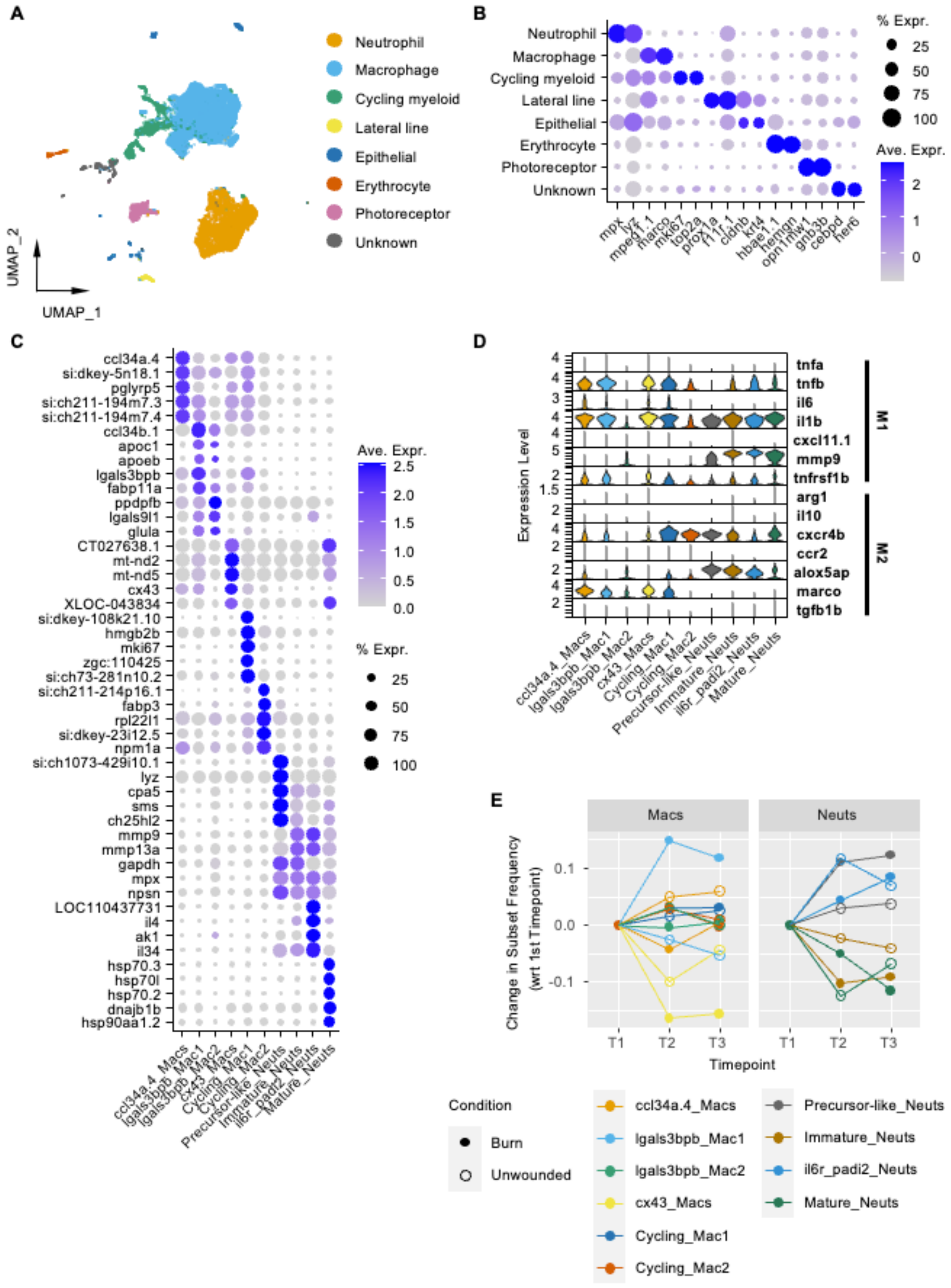
A
Neutrophil
Macrophage
Cycling myeloid
Lateral line
Epithelial
Erythrocyte
Photoreceptor
Unknown
UMAP_1
UMAP_2
B
% Expr.
Ave. Expr.
mpx
lyz
mpeg1.1
marco
mki67
top2a
prox1a
f11r.1
cldnb
krt4
hbae1.1
hemgn
opn1mw1
gnb3b
cebpd
her6
C
Ave. Expr.
% Expr.
ccl34a.4_Macs
lgals3bpb_Mac1
lgals3bpb_Mac2
cx43_Macs
Cycling_Mac1
Cycling_Mac2
Precursor-like_Neuts
Immature_Neuts
il6r_padi2_Neuts
Mature_Neuts
D
Expression Level
tnfa
tnfb
il6
il1b
cxcl11.1
mmp9
tnfrsf1b
arg1
il10
cxcr4b
ccr2
alox5ap
marco
tgfb1b
M1
M2
E
Macs
Neuts
Change in Subset Frequency
(wrt 1st Timepoint)
Timepoint
T1
T2
T3
Condition
Burn
Unwounded

**Supplemental Figure 1 Myeloid cell identity confirmation.** (A) All-cell-included clustering visualized on UMAP axes. (B) Relative expression level of marker genes for each major cell type labeled in (A). Color represents level of expression, while size represents percentage of cells with detectable expression. (C) Relative expression level of top ranking differentially expressed genes in each myeloid subset. Color represents level of expression, while size represents percentage of cells with detectable expression. (D) Expression distribution of zebrafish M1/M2 markers in myeloid subsets. (E) Relative frequency at later time points for each myeloid subsets compared to T1.

Supp. Fig. 2

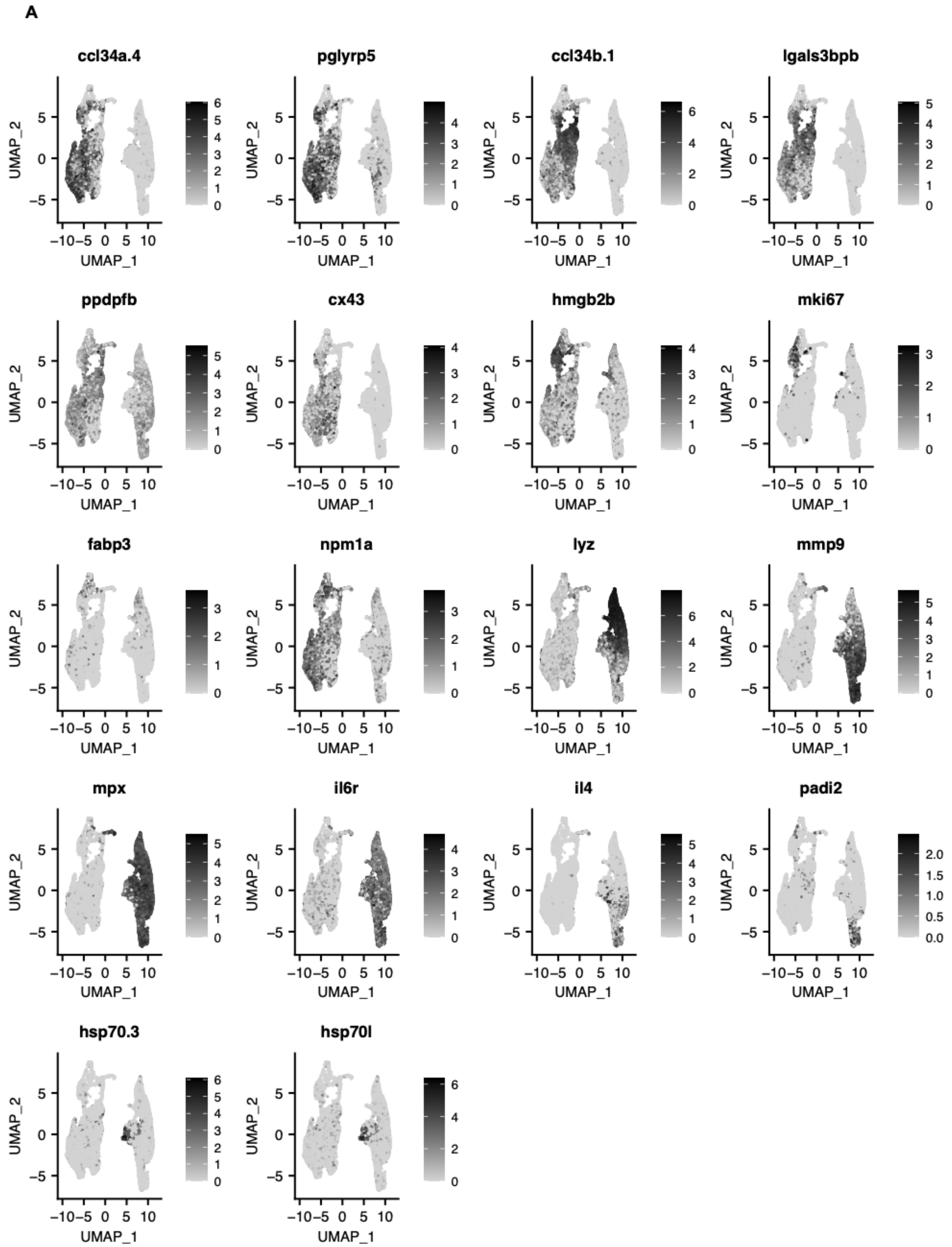


**Supplemental Figure 2 Representative expression distribution of myeloid subset signatures**

**on UMAP axes.** The UMAP axes in use are the same as in Figure 1B, with cells colored by log-normalized expression level.

Supp. Fig. 3

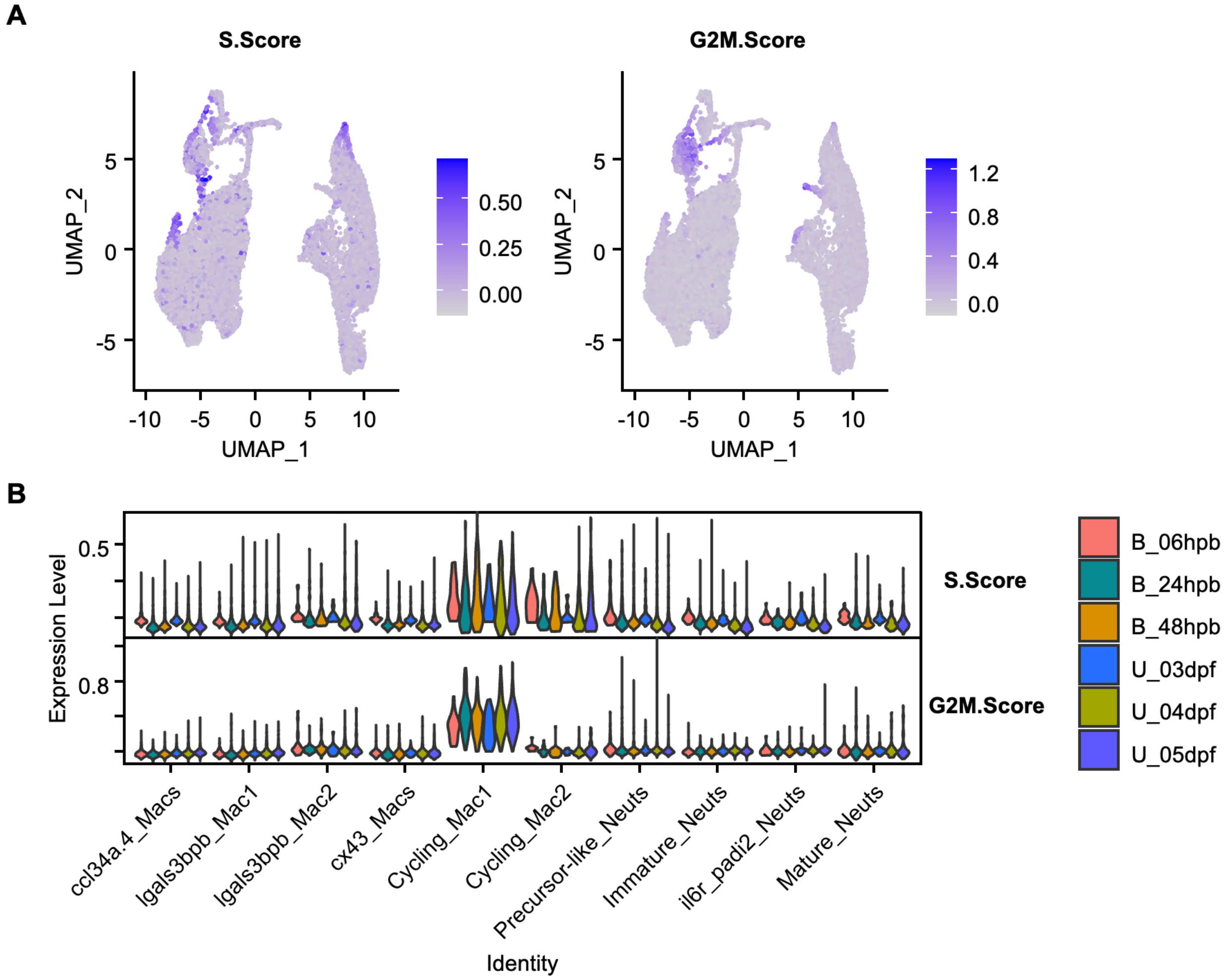


**Supplemental Figure 3 Cell cycle features in myeloid subsets.** (A) Distribution of S and G2M-phase score in myeloid subsets projected onto the same UMAP axes as in Figure 1B. (B) Distribution of S and G2M-phase score in myeloid subsets by condition and time.

Supp. Fig. 4

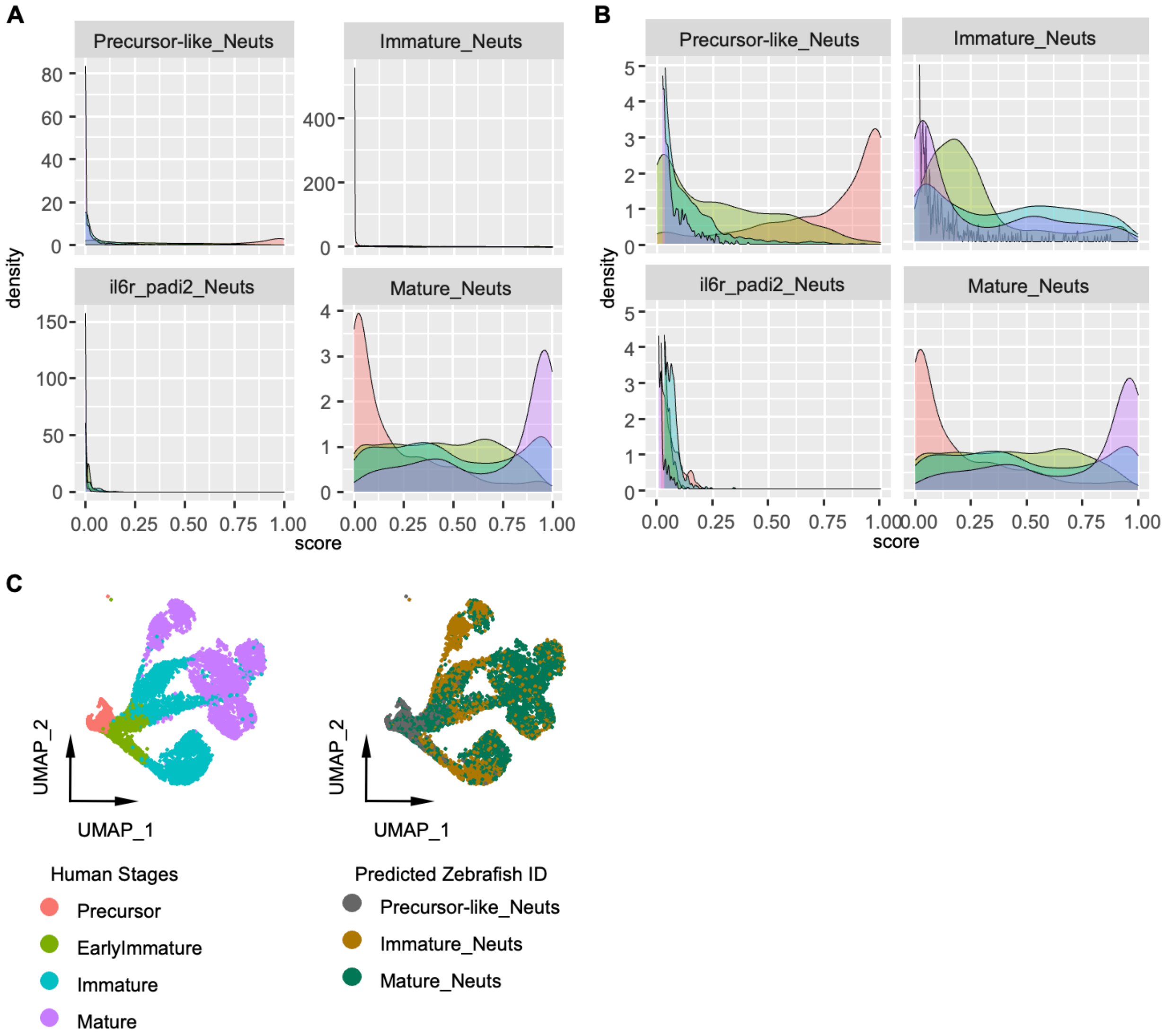


**Supplemental Figure 4 Identity label transfer between human neutrophil stages and zebrafish neutrophil subsets**. (A-B) Distribution of predicted zebrafish subset scores for human neutrophil stages shown as density plots with color codes for human neutrophil stages shared with (C). The y-axis scale was free in (A) while kept at (0, 5) in (B) to better observe the right end of the distribution. (C) Known stage and predicted zebrafish neutrophil subset identity on human neutrophils. UMAP axes are the same as published (6).

Supp. Fig. 5

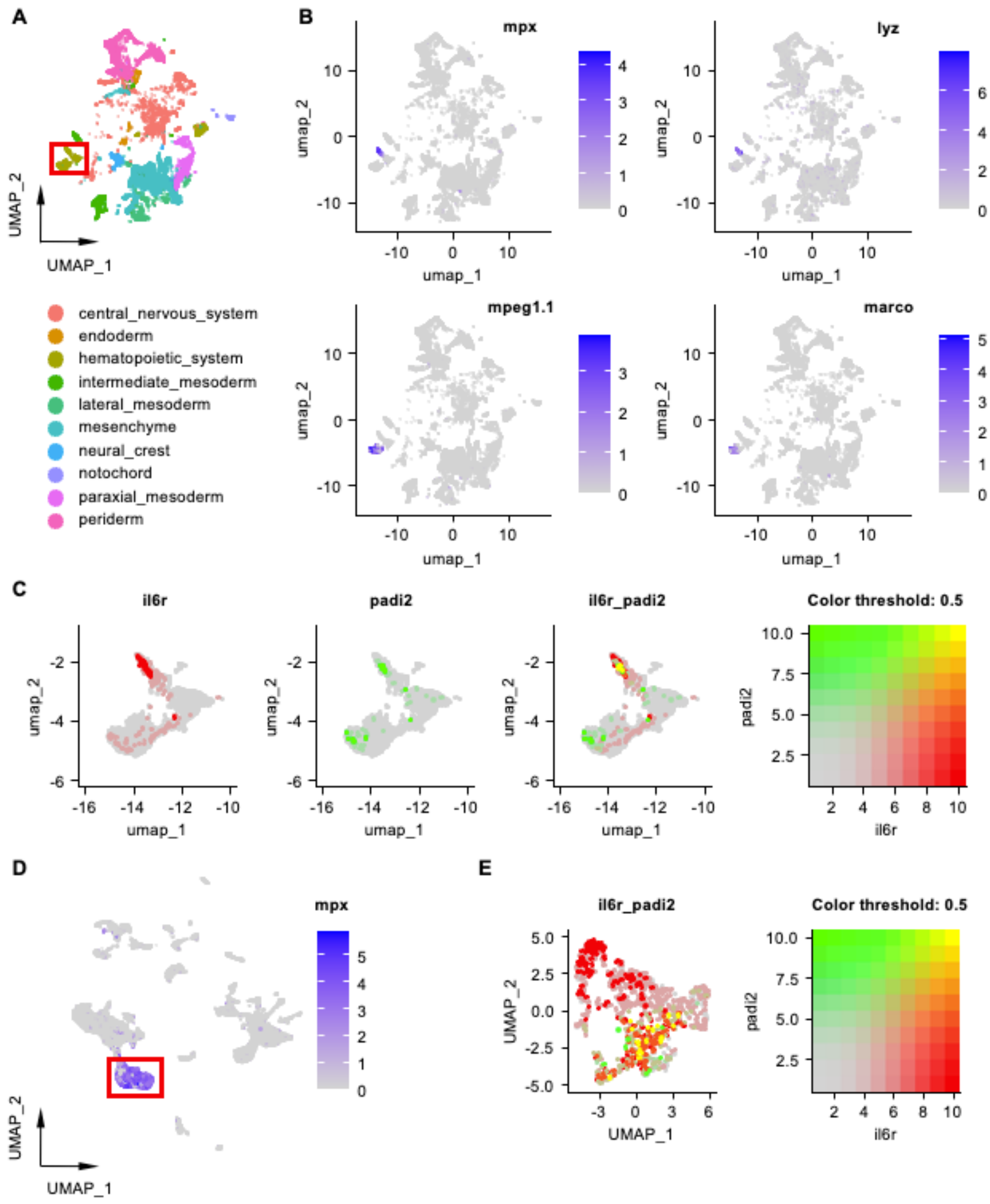


**Supplemental Figure 5 Confirm il6r_padi2_Neuts in ZebraHub and tessellated lymphoid network.** (A-C) Visualization using ZebraHub data where (A) presents all 5 dpf cells from

ZebraHub transcriptomics colored by major anatomy ontology class on UMAP axes. Myeloid cells from the hematopoietic system cluster highlighted in a red rectangle. (B) Expression level of neutrophil markers (*mpx*, *lyz*) and macrophage markers (*mpeg1.1*, *marco*) in 5 dpf cells visualized on the same UMAP axes as in (A). (C) Co-expression pattern for *il6r* and *padi2* in myeloid cells from all time points. Expression levels of the two genes are colored by the color map on the right. (D-E) Visualization using adult tessellated lymphoid network (TLN) data where (D) presents neutrophil marker gene *mpx* expression in all cells on UMAP axes. Neutrophils cluster is highlighted in a red rectangle. (E) Co-expression pattern for *il6r* and *padi2* in TLN neutrophils. Expression levels of the two genes are colored by the color map on the right.

Supp. Fig. 6

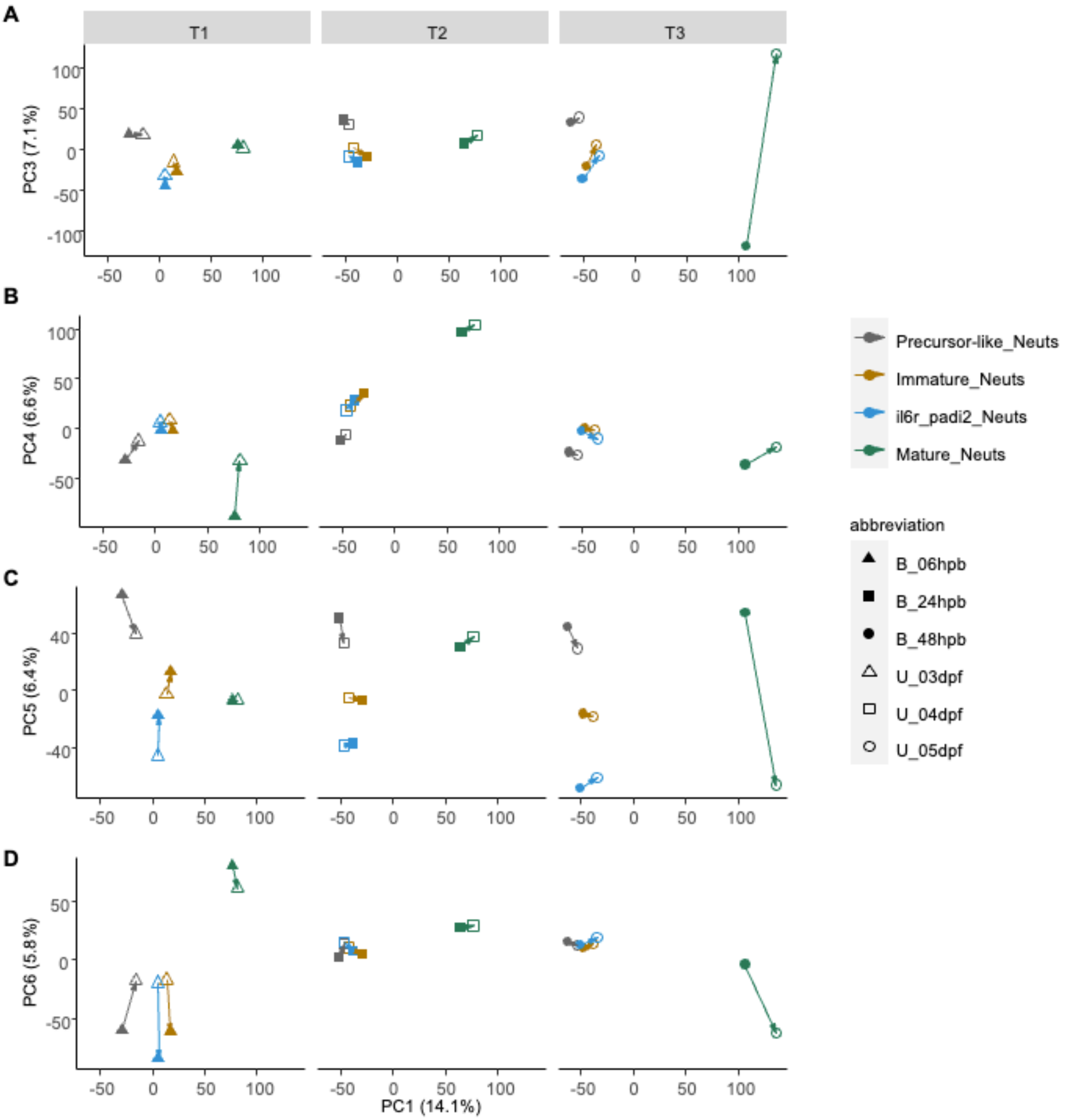


**Supplemental Figure 6 Neutrophil subsets on PC axes.** (A-D) Visualization for neutrophil subset relationship grouped by time between PC1 and (A) PC3, (B) PC4, (C) PC5, (D) PC6.

Supp. Fig. 7

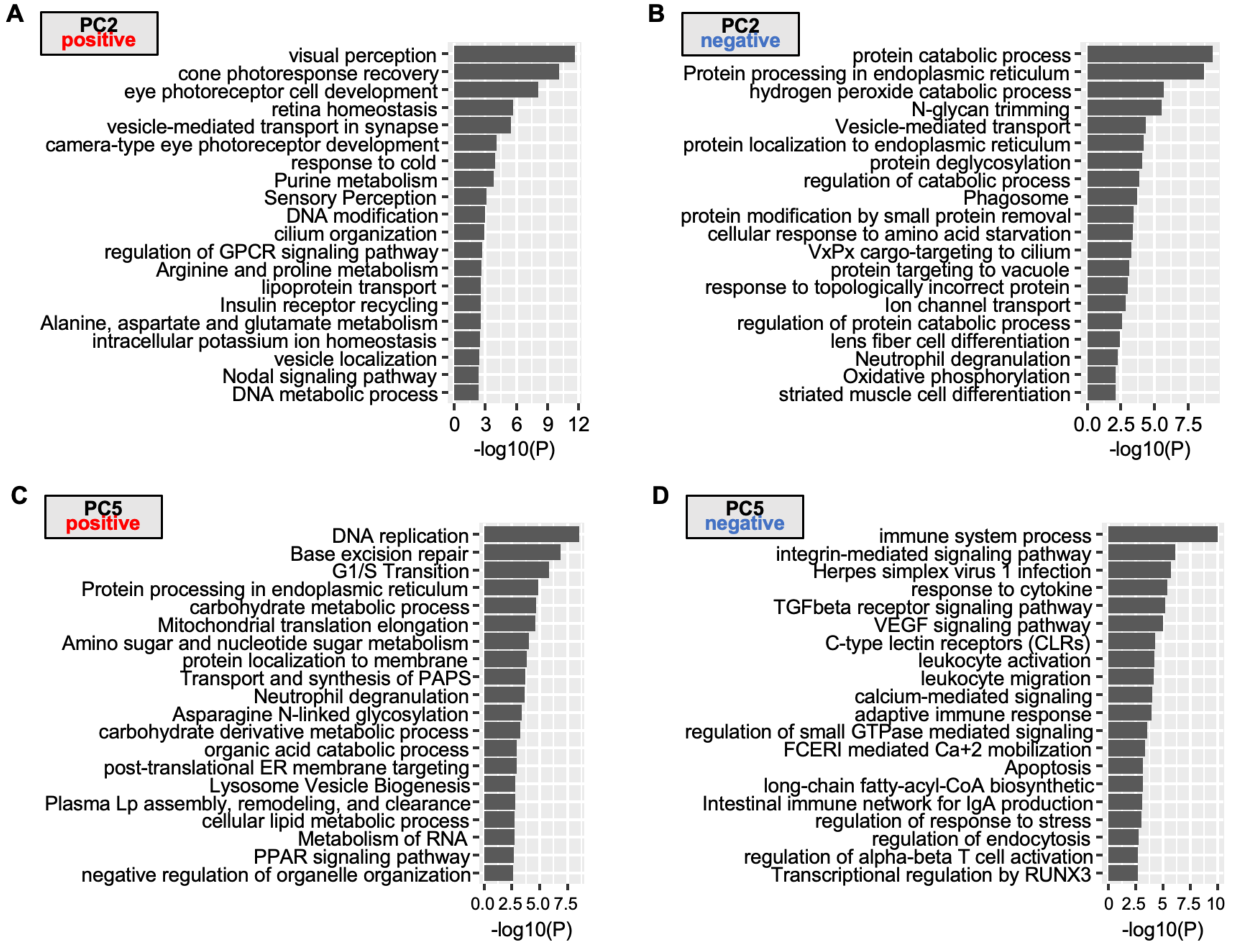


**Supplemental Figure 7 Functional annotation of PC associated genes.** (A-D) Enriched GO terms for genes (A) positively associated with PC2, (B) negatively associated with PC2, (C) positively associated with PC5, (D) negatively associated with PC5.

Supp. Fig. 8

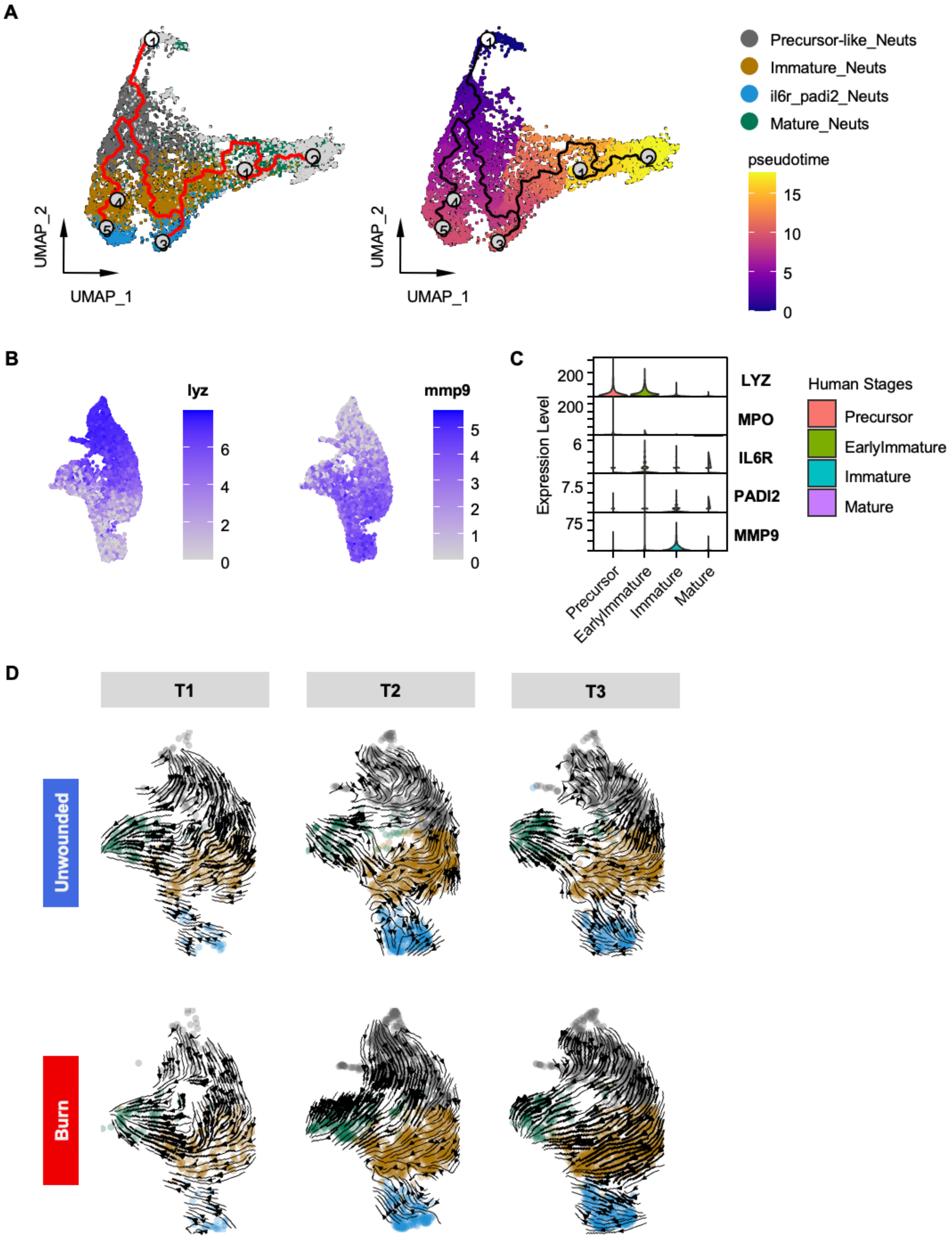


**Supplemental Figure 8 Orthogonal trajectory analysis of neutrophil subsets.** (A) Pseudotime analysis of neutrophils using Monocle 3. UMAP projection independently calculated to show predicted trajectory (left) and pseudotime (right). (B) Expression distribution of markers related to zebrafish neutrophil maturity on UMAP axes that are shared with Figure 3A. (C) Expression level distribution of general neutrophil marker gene and Il6r_padi2_Neuts signatures in human neutrophils. (D)  Velocity prediction for neutrophils all three time points projected as arrows on the same UMAP axes as in Figure 3A.

Supp. Fig. 9

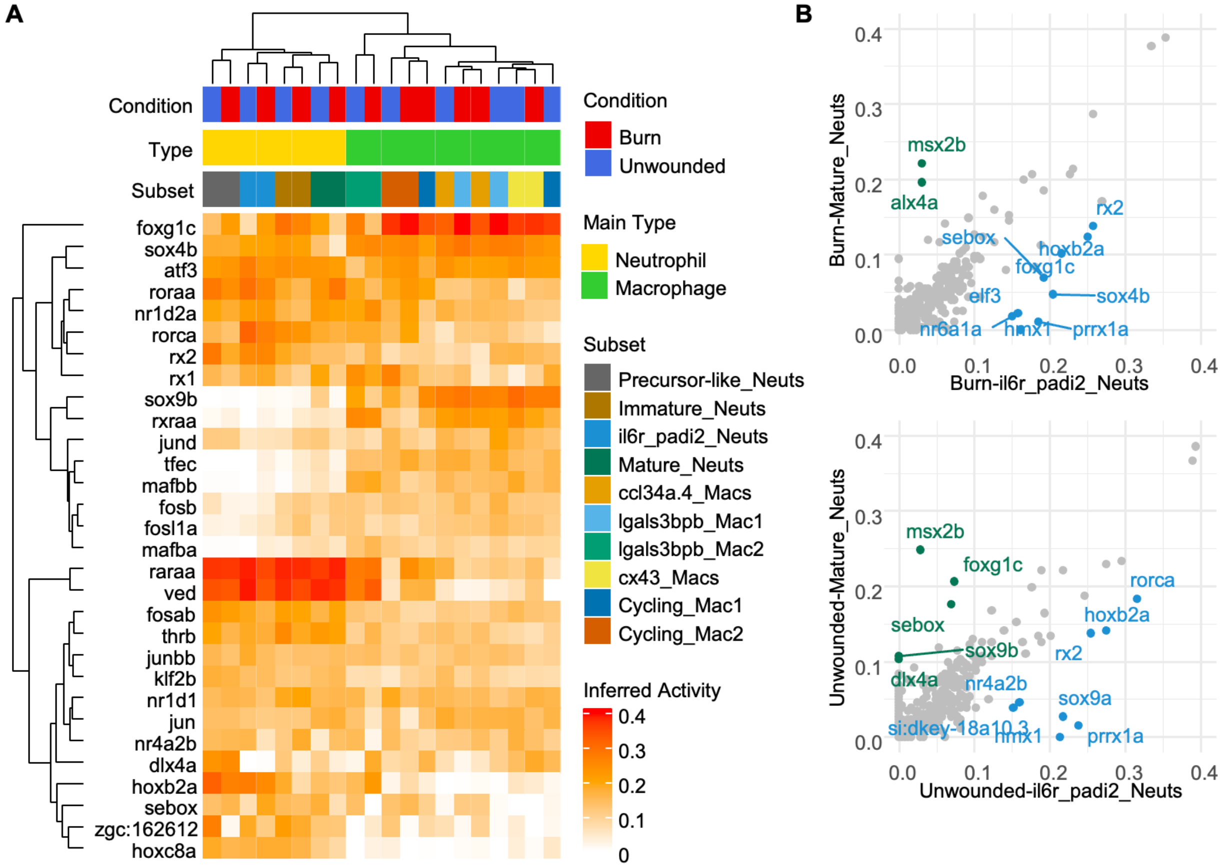


**Supplemental Figure 9 Transcription factor comparison among myeloid subsets.** (A) Degree of centrality of top 30 transcription factors in myeloid subsets. Level of degree of centrality marked as Inferred Activity. (B) Pairwise comparison of TF's degree of centrality between Mature_Neuts and il6r_padi2_Neuts in either burn or unwounded conditions.

Supp. Fig. 10

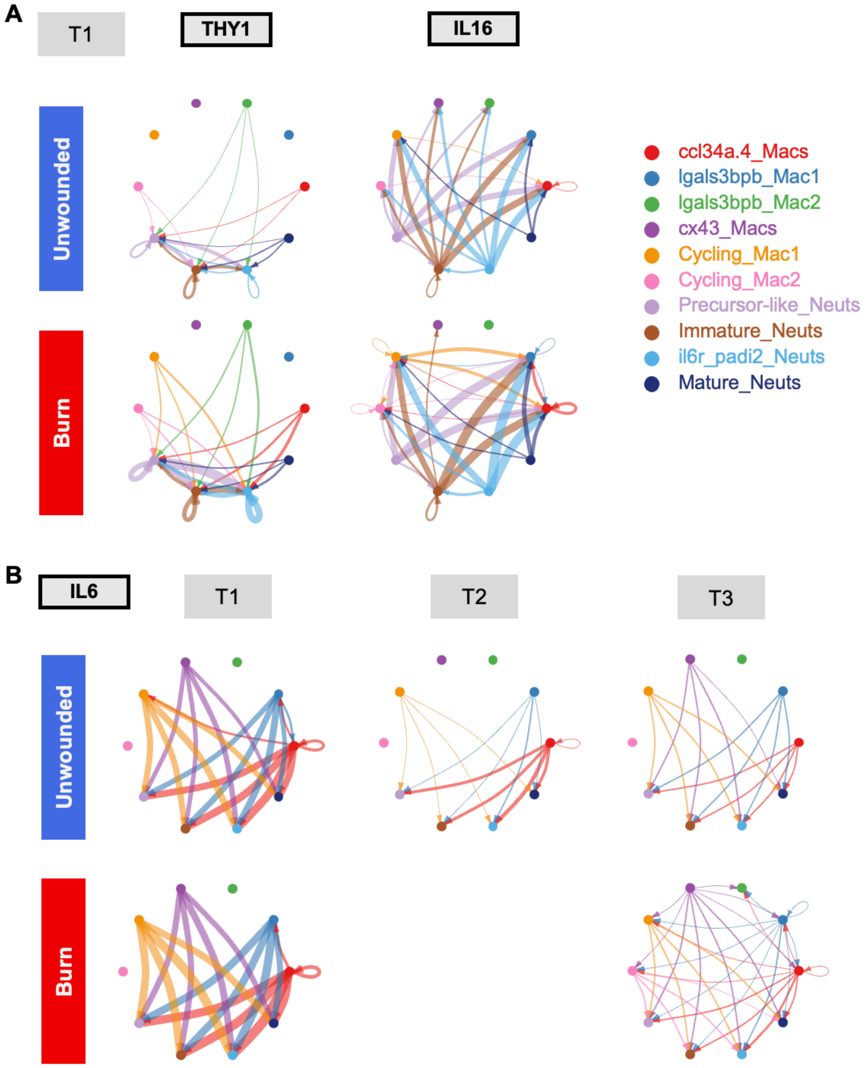


**Supplemental Figure 10 Pathway-specific network.** Signal input and output among myeloid subsets shown as network plot for (A) T1 only for THY1 and IL16 pathway, and (B) all time points for IL6 pathway. Myeloid subset color coding is shared.
